# Adipose Tissue Pericellular Fibrosis, Weight Loss and Features of Metabolic Dysfunction-Associated Steatotic Liver Disease after Bariatric Surgery

**DOI:** 10.64898/2026.10.07.757298

**Authors:** Eve-Julie Tremblay, Juan de Toro Martin, Stéfane Lebel, Christian Y. Couture, Fannie Lajeunesse-Trempe, Ina Maltais-Payette, Giada Ostinelli, Marie-Claude Vohl, Laurent Biertho, Pascale Mauriège, André Tchernof

## Abstract

**Introduction:** Adipose tissue fibrosis has been associated with metabolic dysfunction-associated steatotic liver disease (MASLD) and suboptimal weight loss after bariatric surgery. The objective was to characterize changes in adipose tissue fibrosis and extracellular matrix (ECM) components in adipose tissues from two abdominal fat compartments, subcutaneous and omental, in response to bariatric surgery; and determine the effect of weight loss on the association between adipose tissue fibrosis and MASLD feature changes with weight loss.

**Methods:** Liver and adipose tissue biopsy samples were obtained before and after weight loss from patients who underwent two-stage bariatric surgery. Adipose tissue pericellular fibrosis was evaluated histologically with picrosirius red staining for collagen quantification and normalized to adipocyte count per microscope field. Transcript levels of ECM remodeling and inflammatory genes were measured by RNA sequencing in subcutaneous adipose tissue pre- and post-surgery.

**Results:** At baseline, adipose tissue pericellular fibrosis did not associate with MASLD features, metabolic dysfunction-associated steatohepatitis or hepatic fibrosis in patients with severe obesity. Changes in liver histology after surgery were unrelated to changes in adipose tissue fibrosis. Transcripts for most genes involved in ECM remodeling and inflammation were downregulated in subcutaneous fat one year after bariatric surgery. Omental, but not subcutaneous, adipose tissue fibrosis decreased after surgery when adjusted for cell number (mean ± SD: 0.41±0.12 vs 0.32±0.08 µm²/cell, p<0.005). Higher subcutaneous adipose tissue fibrosis correlated with reduced weight loss (estimate ± SE: −2.04 ± 0.99, p<0.05) and BMI loss (−0.73 ± 0.35, p<0.05), while increased omental adipose tissue fibrosis after weight loss was associated with worsening hepatic fibrosis (0.31 ± 0.14, p<0.05).

**Conclusion:** In individuals with severe obesity undergoing bariatric surgery, adipose tissue fibrosis is poorly correlated with MASLD features. Notably, pericellular fibrosis in subcutaneous abdominal, but not omental, adipose tissue was associated with a reduced weight-loss response to surgery.

## INTRODUCTION

Clinical obesity is a chronic disease in which functional alterations of tissues and organs are caused by excess adiposity (Rubino et al., 2025). It represents an important risk factor for several chronic diseases such as arterial hypertension, type 2 diabetes mellitus, dyslipidemia, metabolic dysfunction-associated steatotic liver disease (MASLD), and coronary heart disease (Wharton et al., 2020). Individuals living with severe obesity [body mass index (BMI) ≥ 40 kg/m^2^] are at even higher risk of co-morbidities, mortality, and reduced life expectancy (GBD 2015 Obesity Collaborators et al., 2017). Weight loss induced through metabolic bariatric surgery is the most effective treatment for eligible patients, leading to durable remission of many obesity-related comorbidities (Arterburn et al., 2020). However, the response to bariatric surgery is subject to large inter-individual variability, with some patients experiencing suboptimal weight loss or weight regain (Noria et al., 2023).

Adequate adipose tissue extracellular matrix (ECM) remodeling has been proposed as a key factor for achieving and maintaining successful weight loss after bariatric surgery (Gliniak et al., 2023). The ECM is a dynamic structure that is constantly remodelled to offer physical support to adipocytes; as such, ECM remodeling is a process allowing adipose tissue expansion (Ruiz-Ojeda et al., 2019). However, excess adiposity, and most specifically, unresolved, low-grade inflammation, leads to abnormal collagen deposition and fibrosis development that is tightly associated with macrophage infiltration (Reggio et al., 2013; Sun et al., 2011). This excess of ECM components impairs adipose tissue mass expansion and worsens adipose tissue dysfunction, thereby contributing to metabolic dysregulation (Khan et al., 2009). Subcutaneous abdominal (SCABD) adipose tissue fibrosis measured in severe obesity is associated with resistance to weight loss induced by bariatric surgery (Abdennour et al., 2014). Gastric bypass surgery is also less effective in patients with high amounts of SCABD adipose tissue fibrosis (Divoux et al., 2010). In other studies, although fibrosis in SCABD or visceral adipose tissue prior to surgery did not correlate with weight loss after Roux-en-Y gastric bypass or sleeve gastrectomy, collagen Vα1 (*COL5A1*) mRNA levels in subcutaneous fat were an independent predictor of total weight loss 12 months post-operation (Osorio-Conles, Olbeyra, et al., 2023). The association between subcutaneous fibrosis and weight loss seems well-established, but little is known about the effect of omental fibrosis on weight loss.

On the other hand, bariatric surgery-induced weight loss seems to have limited impact on the amount of fibrosis in fat tissues. An increase in SCABD fat collagen was observed in patients undergoing bariatric surgery one year after the intervention (Liu et al., 2016). Similarly, six months after sleeve gastrectomy, total fibrosis in SCABD adipose tissue of patients was not resolved despite significant weight loss (Chabot et al., 2017). This observation may originate from the method used to measure fibrosis in adipose tissue. Histological measurements of fibrosis, consisting of calculating the ratio of fibrous tissue area stained to total tissue area, is clearly influenced by adipocyte size. Because adipocytes are smaller after weight loss (Murphy et al., 2017), they leave more space for collagen deposits to be colored per surface area. The stained areas of collagen may then be overestimated after weight loss, which may lead to the false conclusion that adipose fibrosis increases after weight loss.

Studies on adipose tissue fibrosis have also shown a correlation with features of MASLD and insulin resistance. For example, SCABD fat fibrosis was higher in individuals with obesity and MASLD than in those with obesity alone and inversely correlated with both hepatic and whole-body insulin sensitivity (Beals et al., 2021). Yet, in women with abdominal obesity, visceral but not SCABD adipose tissue pericellular fibrosis correlated with higher fasting insulin levels (Michaud et al., 2016). Another study showed no correlation between omental (OM) adipose tissue fibrosis and liver fibrosis in bariatric surgery candidates but did report an association with SCABD fat fibrosis (Abdennour et al., 2014). This suggests that the association between adipose tissue fibrosis, MASLD and insulin resistance may be depot-specific. The impact of weight loss on this association also remains to be determined.

The objectives of this study were to examine the effect of bariatric surgery on changes in adipose tissue fibrosis and ECM components from two fat compartments, SCABD and OM adipose tissues, in response to bariatric surgery, and to determine whether excess adipose tissue fibrosis associates with limited weight loss. Additionally, the effect of bariatric surgery on the association between adipose tissue pericellular fibrosis and MASLD feature changes were evaluated. Finally, measurements of adipose tissue collagen by picrosirius red staining expressed per surface area or per cell number were compared to account for weight loss-induced changes in adipocyte size.

## PARTICIPANTS AND METHODS

Patients meeting the criteria for bariatric surgery (BMI ≥ 40 kg/m^2^ or BMI ≥ 35 kg/m^2^ with at least one diagnosed comorbidity including type 2 diabetes mellitus, arterial hypertension, dyslipidemia, or sleep apnea) underwent bariatric surgery at the Quebec Heart and Lung Institute (QHLI), in Canada. Patient selection for surgery followed the standard National Institutes of Health recommendations (Hubbard & Hall, 1991). Exclusion criteria were general contra-indications to bariatric surgery, a BMI<35 kg/m^2^, age under 18 or over 60 years, abnormal bowel habits including chronic inflammatory bowel disease, pregnancy, cirrhosis or albumin deficiency. All patients provided written informed consent before being included in the study.

### Patient samples 1 and 2

For cross-sectional analyses (sample 1), plasma, SCABD and OM adipose tissues, as well as liver samples from 269 patients were retrieved from the QHLI Obesity Biobank, in accordance with institutionally approved management modalities. For pre- and post-weight loss analyses of SCABD and OM adipose tissues as well as liver tissues (sample 2), 38 other patients underwent a two-stage bariatric surgery at the QHLI. This consists of a sleeve gastrectomy followed by a biliopancreatic diversion with duodenal switch once sufficient weight loss has been achieved as a result of the gastrectomy. Plasma, SCABD and OM adipose tissues as well as liver samples were obtained at the time of the first surgery, and then 26 ± 6 months later on average, during the second surgery. The experimental protocol was approved by the ethics committee of the QHLI-Université Laval (approval numbers #2017-2710, #2018-2951 and #2021–3398).

SCABD and OM adipose tissues were collected at the site of trocar insertion (lower abdomen) and from the proximal portion of the greater omentum, respectively. A portion of the sample was fixed in 10% formalin and then processed for standard paraffin embedding. Five-micrometer slices of SCABD and OM adipose tissues were mounted on the same slide for each patient. The experimental protocol was approved by the ethics committee of the QHLI-Laval University (approval numbers #2021–3398 and #2018-2951).

### Patient sample 3

For prospective analyses on SCABD samples, a total of 21 patients received one of three surgeries performed at the QHLI (sleeve gastrectomy, Roux-en-Y gastric bypass, or biliopancreatic diversion with duodenal switch). At the time of surgery SCABD adipose tissue samples were surgically removed from each patient. Twelve months later, SCABD adipose tissue was obtained with a surgical biopsy performed under local anesthesia at a site where a trocar was inserted a year before during surgery. The experimental protocol was approved by the ethics committee of the QHLI-Université Laval (approval number #2015-2466). This sample was part of the clinical trial REMISSION which registered at Clinicaltrials.gov (NCT02390973).

### Pericellular fibrosis measurements

These measurements were performed in patient samples 1 and 2. Adipose tissue slides were stained with picrosirius red for collagen quantification (Divoux et al., 2010; Henegar et al., 2008). Our group previously determined that collagen deposition around the adipocyte, especially in the OM fat compartment, is related to total and regional adiposity as well as altered cardiometabolic risk profile, as opposed to whole tissue-based assessments of adipose tissue fibrosis (Michaud et al., 2016). Therefore, collagen surrounding adipocytes (i.e. pericellular collagen) was chosen as a relevant marker for pathological fibrosis accumulation in the study samples. Adipose tissue fibrosis was expressed as the ratio of the sum of stained areas to the sum of field surfaces measured in 10 random fields examined at 10x magnification (Divoux et al., 2010) using image analysis automated with imaging software (ImageJ). Every 10 random field displaying adipocytes and stained collagen was also divided by the number of adipocytes present on each slide at 10x magnification, resulting in the measurement of fibrosis area per cell number in µm^2^/cell.

### RNA sequencing and transcript selection

RNA sequencing of SCABD tissues was performed only in patient sample 3. Transcript abundance was measured as previously described (Bouchard-Mercier et al., 2022). After RNA extraction, library preparation was carried out using the Illumina NEB stranded mRNA library preparation kit (Illumina, San Diego, CA, US) and sequencing was performed on the Illumina NovaSeq 6000 S4 platform (Illumina) using 100 bp paired-end reads. After quality control, read quantification was performed using kallisto (v0.46.2) (Bray et al., 2016) with 100 bootstraps. Reads were further aligned to the GRCh38 human reference transcriptome. The obtained transcript counts were used to infer gene-level abundance estimates with the tximport (v1.20.0) R package (Soneson et al., 2015). Gene expression was normalized, and low expression genes were filtered out with the filterByExpr function in edgeR (v3.34.0) (Robinson et al., 2010).

A total of 18 genes were included in the study (**Suppl Table 1**). Genes associated with ECM components such as collagen [*COL5A1* and collagen Viα3 (*COL6A3*)], and collagen stability (lysyl oxidase; *LOX*, *LOX1* and *LOX2*) were selected (Khan et al., 2009; Mak et al., 2016; Pasarica et al., 2009; Pastel et al., 2018). Genes coding for enzymes regulating the remodeling of the ECM in adipose tissue including metalloproteinases (*MMP2*, *MMP9* and *MMP14*) and metalloproteinase inhibitor (*TIMP2*) were also selected (Berg et al., 2014; Sachan et al., 2023). Other genes related to fibrogenesis such as transforming growth factor beta 1 (*TGFB1*), a regulator of fibrosis (Fain et al., 2005; Meng et al., 2016), platelet-derived growth factor subunit A (*PDGFA*), whose activity drives adipose tissue fibrosis (Marcelin et al., 2017), connective tissue growth factor (*CCN2*), a strong pro-fibrotic factor (Yoshino et al., 2019), and endothelial plasminogen activator inhibitor (*SERPINE1*), a fibrinolysis inhibitor (Kaji, 2016) were also included. Non structural protein-coding gene secreted protein acidic and rich in cysteine (*SPARC*) was selected because the encoded protein regulates cell growth through interactions with the ECM (Kos & Wilding, 2010). Other transcripts were chosen for the association of the encoded protein with inflammation, including macrophage markers cluster of differentiation 68 (*CD68*) and mannose receptor C-type 1 (*MRC1*) (Engin, 2017; Moreno-Navarrete et al., 2013), transcription factor hypoxia-inducible factor 1-alpha (*HIF1A*), whose activity mediates the effects of hypoxia (Halberg et al., 2009; Sun et al., 2013), and inflammasome NOD-like receptor family pyrin domain containing 3 (*NLRP3*) (Pahwa et al., 2021).

### Liver sampling and histological evaluation

Liver biopsies performed as a routine part of all bariatric surgery procedures were taken from the edge of the left liver lobe, fixed in formalin, and stained with hematoxylin/eosin and Masson’s trichrome. The histologic characteristics, including hepatic fibrosis, MASLD and metabolic dysfunction-associated steatohepatitis (MASH) features were rated by trained pathologists according to the Brunt grading system (Brunt et al., 1999).

Patients who underwent two-stage bariatric surgery (patient sample 2) were then classified according to the change in steatosis, MASH and hepatic fibrosis grade. Patients starting with grade 0 and keeping that same grade after surgery were in the “Maintained” group. Patients with grade 1 or more before surgery that improved by at least one grade with bariatric surgery were in the “Improved” group. Patient starting with grade 1 or more who kept the same grade after surgery were in the “Unimproved” group. Patient with grade 1 or more before surgery with worsening of at least one grade after weight loss were in the “Worsened” group. Finally, patients without any steatosis, MASH or fibrosis before surgery, but who worsened by at least one grade after surgery were in the “*de novo*” group. The “Maintained” and “Improved” groups were combined for analysis and compared with the combined “Unimproved”, “Worsened” or “*de novo*” group.

### Statistical analyses

Statistical analyses were performed using JMP software (SAS Institute, Carry, NC, USA). Mean values, percentage, range and SD were used to describe patient sample 1 characteristics. Patient characteristics before and after bariatric surgery were compared using Student *t*-tests and Chi-squared tests in patient samples 2 and 3. Two-sided p<0.05 was considered statistically significant. To investigate the associations between adipose tissue fibrosis and liver characteristics associated with MASLD, ANOVA and *post hoc* analyses were performed (Tukey). For analyses regarding the presence or absence of MASH, pre- and post-surgery comparisons, as well as mRNA gene expression, Student *t*-tests were used. Centered-log ratio was used as the unit for mRNA gene expression. Changes in mRNA gene expression and adipose tissue fibrosis following bariatric surgery were correlated to changes in anthropometric measurements, lipid profile and glycemic control by linear regression using delta values. P-values associated with mRNA expression data were adjusted for age and sex, as well as false discovery rate (FDR) using Benjamini-Hochberg. For changes in adipose tissue fibrosis and hepatic characteristics before and after bariatric surgery, an ordinal logistic regression was performed. Patients in the “maintained/improved” group were compared to those in the “unimproved/worsened/de novo” group according to steatosis and hepatic fibrosis grade, as well as the presence or absence of MASH using linear mixed-effects models. Fixed effects were time, group, time × group, and subject ID was a random effect to account for within-subject correlations. *Post hoc* analyses were based on Tukey-Kramer p values.

## RESULTS

Characteristics of the patient samples are presented in **Table 1**. As expected, in addition to reductions in body weight, BMI and waist circumference, patients who underwent bariatric surgery showed significant improvements in their plasma lipid profile insulin-glucose homeostasis. Percent steatosis decreased markedly, from 29.5 ± 26.8% to 7.4 ± 10.0% on average (p<0.01). In patients from sample 3, steatohepatitis improved in 13 out of 36 patients and worsened in 3, whereas liver fibrosis improved in 12 patients and worsened in 6 (**Suppl Tables 2** and **3**). Patients with worsened hepatic characteristics had higher pre-surgery fasting insulin and HbA1c, as well as higher WC and lower HDL-C after surgery, but similar weight loss compared to patients with improved hepatic characteristics (data not shown).

**Table 1:** Characteristics of patients with severe obesity before and after bariatric surgery.

|  | Sample 1 | Sample 2 |  |  | Sample 3 |  |  |
| --- | --- | --- | --- | --- | --- | --- | --- |
| | Mean ± SD | Pre-op<br>Mean ± SD | 12 months<br>post-op<br>Mean ± SD | p-value | First surgery<br>Mean ± SD | Second surgery<br>(26 ± 6 months)<br>Mean ± SD | p-value |
| Men/Women | 84/185 | 10/11 | - | - | 15/23 | - | - |
| Age (years) | 40.0 ± 10.2 | 52.1 ± 8.0 | 53.1 ± 8.0 | 0.690 | 45.1 ± 10.8 | 47.3 ± 10.7 | 0.362 |
| Surgery |  |  |  |  |  |  |  |
| Sleeve Gastrectomy, n (%) | - | 9 (42.9) | - | - | 38 (100) | - | - |
| RYGB, n (%) | - | 5 (23.8) | - | - | 0 | - | - |
| BPD-DS, n (%) | - | 7 (33.3) | - | - | 0 | - | - |
| **Anthropometrics** |  |  |  |  |  |  |  |
| Weight (kg) | 134.8 ± 23.1 | 124.8 ± 19.6 | 83.1 ± 13.3 | <0.001 | 146.0 ± 26.2 | 116.4 ± 22.1 | <0.001 |
| BMI (kg/m<sup>2</sup>) | 49.8 ± 7.5 | 44.4 ± 6.1 | 29.6 ± 4.1 | <0.001 | 52.6 ± 7.3 | 42.0 ± 6.2 | <0.001 |
| WC (cm) | 138.4 ± 16.5 | 138.7 ± 11.8 | 104.9 ± 10.7 | <0.001 | 144.4 ± 16.0 | 125.6 ± 17.0 | <0.001 |
| Total Weight Loss (%) | - | - | 32.7 ± 8.9 | - | - | 20.0 ± 7.2 | - |
| **Plasma Lipid Profile** |  |  |  |  |  |  |  |
| Triacylglycerols (mmol/L) | 1.69 ± 0.88 | 1.69 ± 0.85 | 1.24 ± 0.57 | 0.050 | 1.77 ± 0.87 | 1.34 ± 0.46 | 0.009 |
| HDL-C (mmol/L) | 1.22 ± 0.30 | 1.14 ± 0.32 | 1.34 ± 0.31 | 0.039 | 1.16 ± 0.26 | 1.37 ± 0.28 | 0.001 |
| LDL-C (mmol/L) | 2.70 ± 0.74 | 2.02 ± 0.86 | 2.33 ± 1.17 | 0.333 | 2.49 ± 0.87 | 2.57 ± 0.76 | 0.679 |
| Total Cholesterol (mmol/L) | 4.67 ± 0.86 | 3.92 ± 1.03 | 4.24 ± 1.29 | 0.386 | 4.44 ± 0.98 | 4.54 ± 0.85 | 0.654 |
| Total Cholesterol to HDL-C ratio | 3.96 ± 0.91 | 3.63 ± 1.07 | 3.41 ± 1.78 | 0.627 | 3.95 ± 1.00 | 3.40 ± 0.75 | 0.009 |
| Apo B (mmol/L) | 0.97 ± 0.42 | 0.86 ± 0.25 | 0.84 ± 0.34 | 0.849 | 0.91 ± 0.24 | 0.91 ± 0.23 | 0.976 |
| **Glucose Homeostasis** |  |  |  |  |  |  |  |
| Fasting Glucose (mmol/L) | 6.0 ± 2.1 | 8.8 ± 2.0 | 5.2 ± 1.2 | <0.001 | 7.2 ± 2.8 | 5.6 ± 1.5 | 0.004 |
| Fasting Insulin (pmol/L) | 177.3 ± 90.0 | 264.0 ± 157.9 | 50.4 ± 24.4 | <0.001 | 268.2 ± 134.8 | 103.5 ± 52.3 | <0.001 |
| HbA1c (%) | 5.7 ± 0.8 | 6.9 ± 1.0 | 5.3 ± 0.6 | <0.001 | 6.6 ± 1.5 | 5.6 ± 0.7 | <0.001 |
| HOMA-IR | 43.0 ± 23.0 | 103.2 ± 65.9 | 11.8 ± 6.5 | <0.001 | 82.5 ± 51.6 | 27.6 ± 21.3 | <0.001 |

SD, Standard Deviation; Min, Minimum; Max, Maximum; BMI, Body Mass Index; WC, Waist Circumference; WHR, Waist-to-Hip Ratio; C, Cholesterol; HDL, High-Density Lipoprotein; LDL, Low-Density Lipoprotein; Apo B, Apolipoprotein B; HbA1c, Glycated Hemoglobin; HOMA-IR, Homeostatic Model Assessment of Insulin Resistance.

### Adipose tissue pericellular fibrosis is not different in MASLD, MASH or hepatic fibrosis in patients with severe obesity

Cross-sectional analyses of patient sample 1 showing pericellular fibrosis in SCABD and OM fat as a function of liver steatosis grade are presented in **Figure 1**. Using the area-based measurement of picrosirius red staining to stain collagen, fibrosis in OM fat was not significantly different across steatosis grades. When expressed relative to adipocyte number, OM fibrosis was higher in patients with steatosis grade S2 compared with those in grade S1. SCABD pericellular fibrosis did not show significant differences as a function of steatosis grade. Patients with MASH showed lower percent of fibrosis in OM fat compared to patients without MASH, but this difference was no longer significant when adipose tissue pericellular fibrosis was expressed relative to cell number (**Figure 2**). Adipose tissue fibrosis did not differ according to liver fibrosis grades (**Figure 3**).

**Figure 1.**
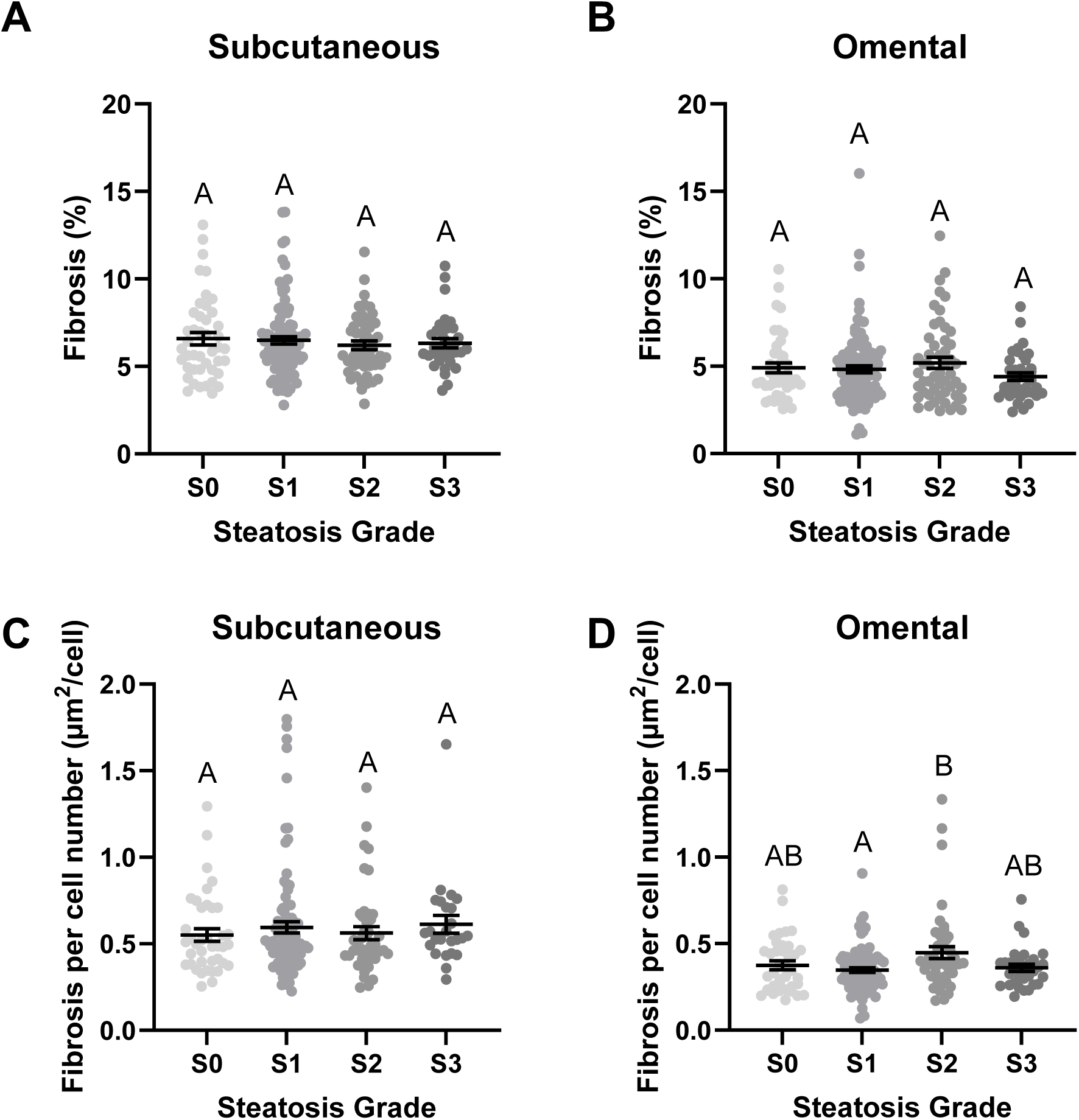
Adipose tissue fibrosis measured by picrosirius red labeling of collagen expressed as area (A, B) or area by cell number (C, D) in two fat compartments according to steatosis grade. P-values are adjusted for age and sex. ANOVA and Tukey, means ± SEM, groups showing different letters are statistically different from each other at p<0.05.

**Figure 2.**
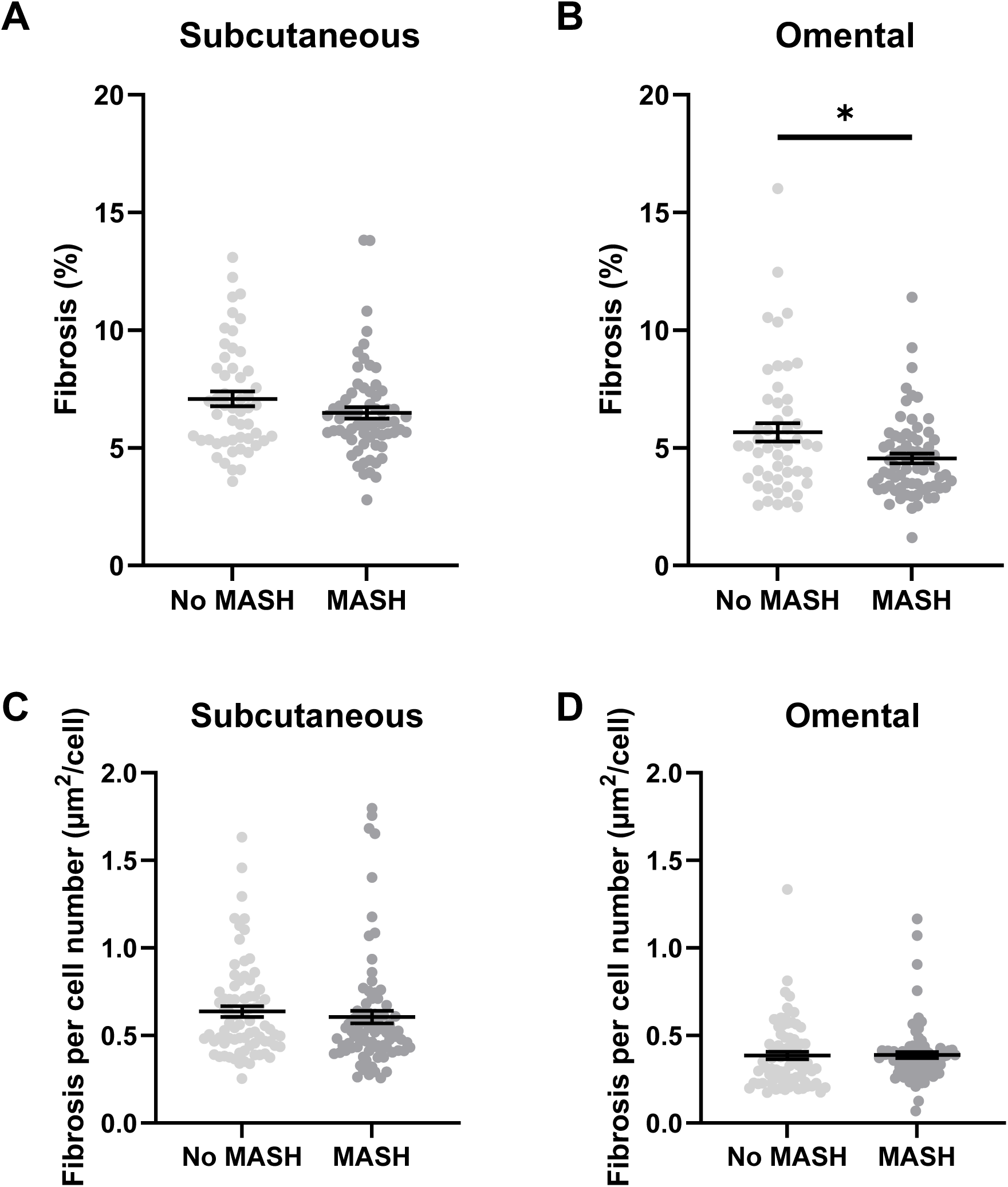
Adipose tissue fibrosis measured by picrosirius red labeling of collagen expressed as area (A, B) or area by cell number (C, D) in two fat compartments according to the presence or absence of MASH. P-values are adjusted for age and sex. Student *t*-tests, means ± SEM, *p<0.05.

**Figure 3.**
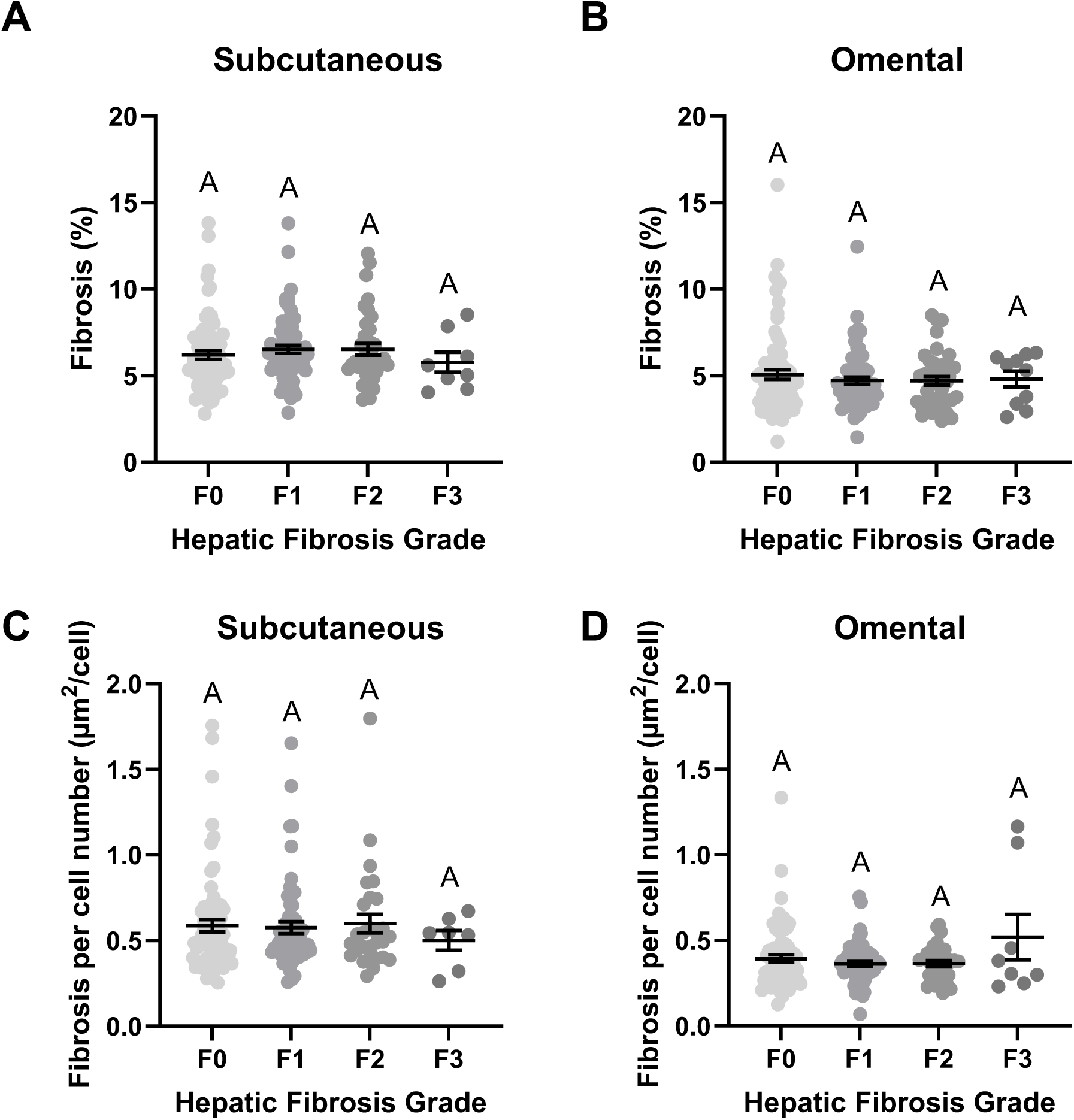
Adipose tissue fibrosis measured by picrosirius red labeling of collagen expressed as area (A, B) or area by cell number (C, D) in two fat compartments according to hepatic fibrosis grade. P-values are adjusted for age and sex. ANOVA and Tukey, means ± SEM, groups showing different letters are statistically different from each other at p<0.05.

### Changes in histological MASLD features after bariatric surgery do not associate with changes in adipose tissue fibrosis

Logistic regression analyses presented in **Table 2** demonstrate that changes in steatosis grade, MASH diagnosis, or hepatic fibrosis following sleeve gastrectomy do not associate with changes in SCABD adipose tissue pericellular fibrosis even when adjusted per cell number. Increases in non-adjusted OM adipose tissue pericellular fibrosis after bariatric surgery were associated with worsening of hepatic fibrosis. To understand whether associations between MASLD and adipose tissue fibrosis differed between responders and non-responders, we grouped patients with improved or worsened steatosis grade, MASH or hepatic fibrosis grade (**Suppl Figures 1**, **2** and **3**). There were no differences in weight-, BMI- or waist circumference-loss between the two groups (data not shown). Patients with improved/unchanged steatosis and MASH had lower adjusted OM adipose tissue fibrosis. Both improved and worsened liver fibrosis groups improved the adjusted amount of OM adipose tissue fibrosis post-surgery. Patients with improved/unchanged liver steatosis also displayed higher levels of unadjusted SCABD fat fibrosis after bariatric surgery.

**Table 2.** Ordinal logistic regression for changes in adipose tissue fibrosis and changes in hepatic characteristics before and after bariatric surgery.

|  | SCABD Fibrosis |  |  | SCABD Fibrosis per Cell Number |  |  |
| --- | --- | --- | --- | --- | --- | --- |
|  | Estimate | Std Err | p-value | Estimate | Std Err | p-value |
| Steatosis Grade | 0.19 | 0.16 | 0.229 | 3.15 | 3.15 | 0.318 |
| MASH | 0.04 | 0.16 | 0.794 | -0.44 | 3.34 | 0.896 |
| Hepatic Fibrosis Grade | -0.22 | 0.14 | 0.124 | -4.27 | 3.09 | 0.166 |
|  | OM Fibrosis |  |  | OM Fibrosis per Cell Number |  |  |
|  | Estimate | Std Err | p-value | Estimate | Std Err | p-value |
| Steatosis Grade | 0.04 | 0.14 | 0.772 | -3.88 | 4.39 | 0.377 |
| MASH | 0.16 | 0.15 | 0.264 | 5.98 | 4.54 | 0.188 |
| Hepatic Fibrosis Grade | **0.31** | **0.14** | **0.028** | 3.03 | 4.00 | 0.449 |

Significant relationships are in bold. MASH, Metabolic dysfunction-associated steatohepatitis. For abbreviations, see Table 1.

### Expression of genes associated with adipose tissue ECM remodeling in SCABD is downregulated after bariatric surgery

Previously published data using this patient sample showed SCABD remodeling after bariatric surgery (Bouchard-Mercier et al., 2022). As shown in **Figure 4**, SCABD adipose tissue mRNA expression of *COL5A1* did not change 12 months post bariatric surgery, and *COL6A3* expression tended to increase after weight loss (FDR-adjusted p=0.07). Likewise, expression of metalloproteinases *MMP2* and *MMP14* did not change after surgery, while expression of *MMP14* and *TIMP2* decreased significantly (both p<0.005). Expression of key regulators of adipose tissue ECM remodeling and inflammation (*SERPINE1*, *CCN2*, *SPARC*, *HIF1A*, *PDGFA*, *TGFB1, MRC1, NLRP3* and *CD68*) all decreased after weight loss (0.005<p<0.01, **Figure 5**). Messenger RNA expression of the aforementioned genes was still significantly lowered after weight loss when type of surgery was examined, with no differences between the three surgery types (data not shown).

**Figure 4.**
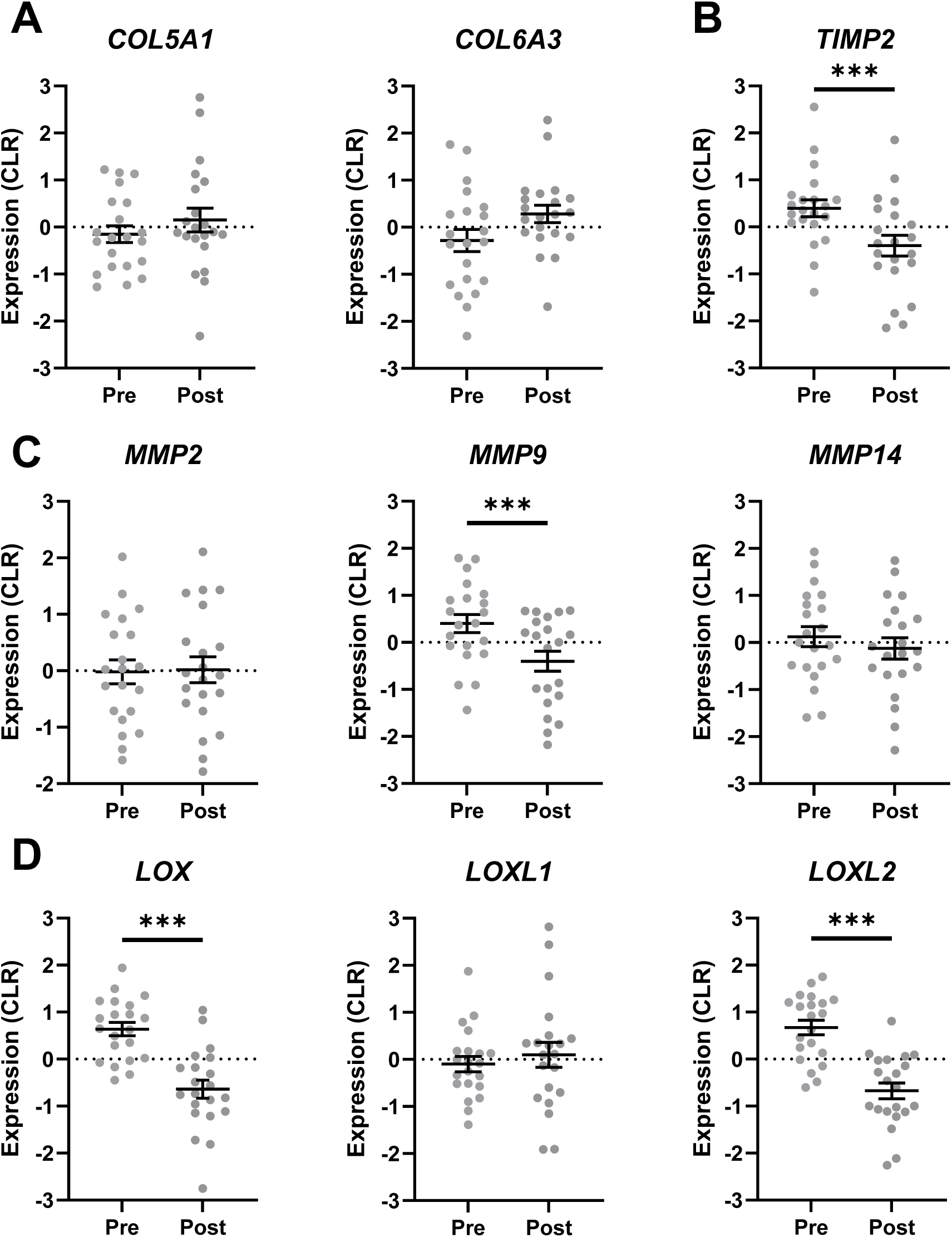
Messenger RNA expression of adipose tissue collagens (A), metalloproteinases (C) and their inhibitor (B), as well as lysyl oxidase (LOX, D) in subcutaneous abdominal adipose tissue before and 12-months after bariatric surgery. P-values are adjusted for age, sex and FDR. Student *t*-tests, means ± SEM, ***p<0.005. CLR, centered-log ratio.

**Figure 5.**
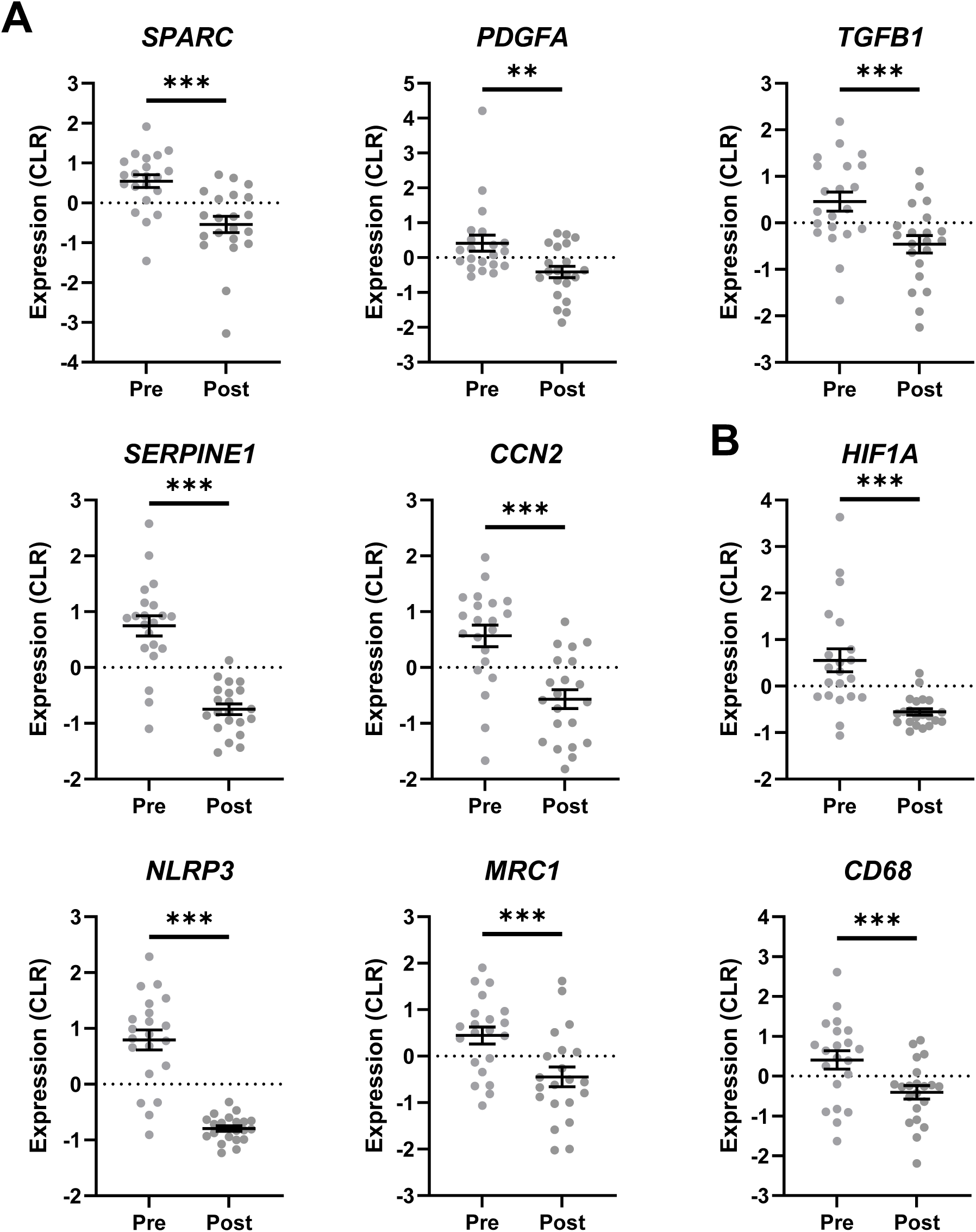
Messenger RNA expression of key regulators of genes coding for extracellular matrix remodeling (A) and inflammation (B) in subcutaneous abdominal adipose tissue before and 12-months after bariatric surgery. P-values are adjusted for age, sex and FDR. Student *t*-tests, means ± SEM, ***p<0.005, ** p<0.01. CLR, centered-log ratio.

Linear regression analyses were used to determine whether changes in mRNA gene expression levels associated with either weight or BMI loss one year after bariatric surgery (**Table 2**). Expression of *COL6A3*, *MMP14* and *MRC1* negatively associated with weight loss, although these associations were no longer significant after FDR adjustment. Conversely, changes in mRNA gene expression of *COL6A3* and *MRC1* mRNA expression were positively associated with BMI loss, however, these associations were no longer significant after FDR correction. No correlations were found between changes in mRNA expression and changes in lipid profile or glycemic profile (data not shown).

### Changes in SCABD adipose tissue pericellular fibrosis associate with weight loss following bariatric surgery

**Figure 6** shows an increase in SCABD adipose tissue pericellular fibrosis after sleeve gastrectomy (p<0.05). When adjusting fibrosis per cell number, changes in SCABD adipose tissue fibrosis were no longer significant. Pericellular fibrosis in OM fat was not different after bariatric surgery but was decreased when adjusted for cell number (p<0.005).

**Figure 6.**
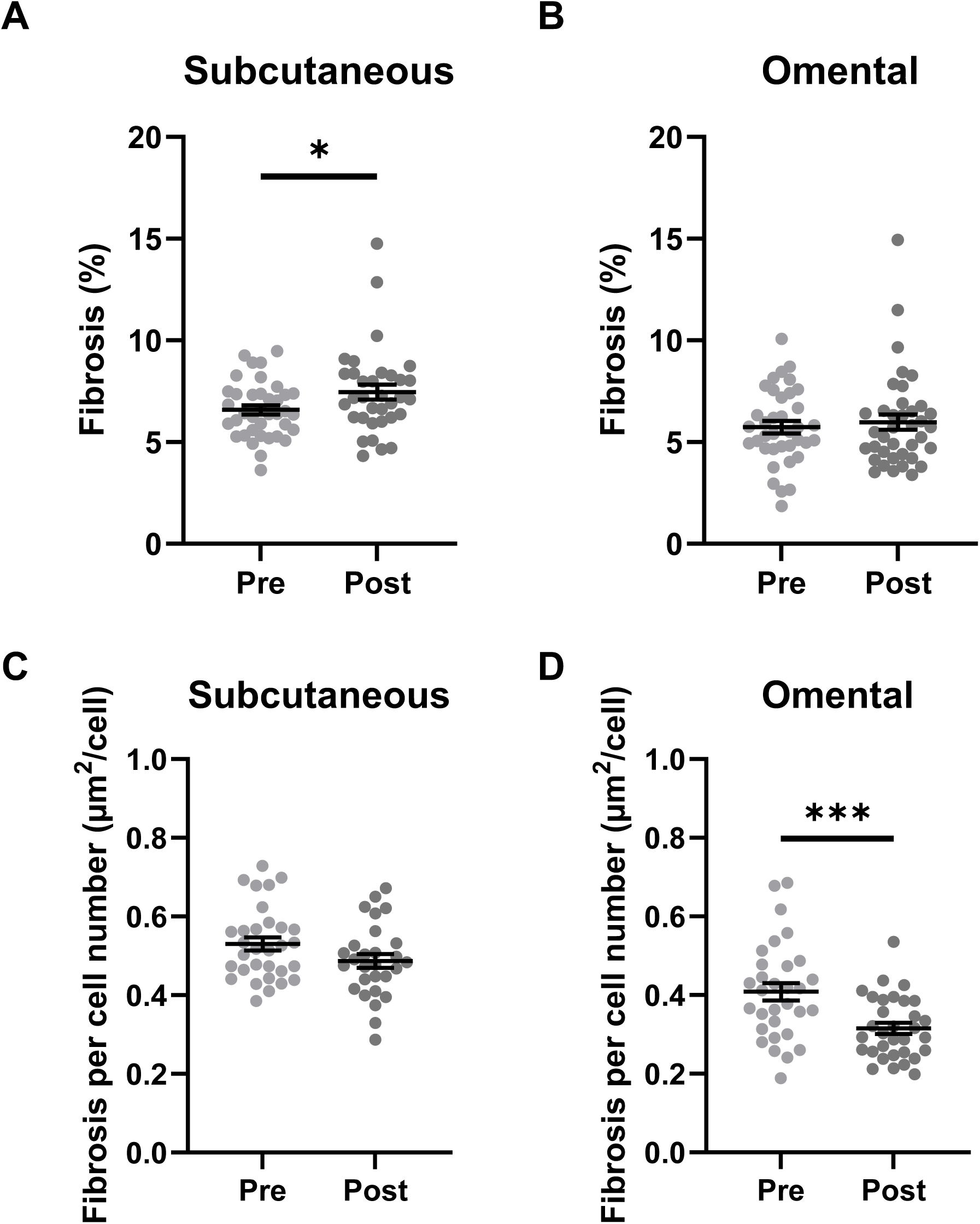
Adipose tissue fibrosis measured by picrosirius red labeling of collagen expressed as area (A, B) or area by cell number (C, D) in two fat compartments before and after bariatric surgery. P-values are adjusted for age and sex. Student *t*-tests, means ± SEM, ***p<0.005, * p<0.05.

We then examined the relationship between pre- and post-surgery adipose tissue pericellular fibrosis and weight loss. There was no correlation between adipose tissue pericellular fibrosis before bariatric surgery and changes in weight, BMI or waist circumference. Post-surgery, OM adipose tissue fibrosis negatively associated with both weight (r=-0.32; p=0.05) and BMI loss (r=-0.42; p=0.008). Total weight loss (%) was divided into tertiles, and pericellular adipose tissue fibrosis was not different between the groups (data not shown). Then, linear regression analyses were performed using either adipose tissue fibrosis area or values adjusted per cell number and anthropometric measurements (**Table 3**). The analysis showed that changes in SCABD fibrosis was negatively associated with changes in weight and BMI, implying that a greater decrease in SCABD fibrosis was associated with a greater decrease in weight and BMI after surgery. No correlations were found between changes in adipose tissue fibrosis and changes in lipid profile or glycemic profile (data not shown).

**Table 3.** Linear regression models for changes in mRNA expression of genes associated with adipose tissue ECM remodeling and changes in anthropometric measurements before and after bariatric surgery.

|  | Weight loss (kg) |  |  |  | BMI loss (kg/m<sup>2</sup>) |  |  |  |
| --- | --- | --- | --- | --- | --- | --- | --- | --- |
|  | Estimate | Std Err | p-value | FDR-adjusted p-value | Estimate | Std Err | p-value | FDR-adjusted p-value |
| *COL5A1* | -0.07 | 0.06 | 0.260 | 0.474 | 0.24 | 0.17 | 0.174 | 0.358 |
| *COL6A3* | **-0.16** | **0.05** | **0.009** | 0.096 | **0.34** | **0.16** | **0.042** | 0.245 |
| *MMP2* | -0.03 | 0.07 | 0.647 | 0.832 | 0.04 | 0.19 | 0.841 | 0.841 |
| *MMP9* | 0.01 | 0.04 | 0.768 | 0.857 | 0.08 | 0.12 | 0.526 | 0.771 |
| *MMP14* | **-0.13** | **0.05** | **0.023** | 0.136 | 0.30 | 0.15 | 0.063 | 0.245 |
| *TIMP2* | -0.03 | 0.06 | 0.629 | 0.832 | 0.08 | 0.16 | 0.620 | 0.771 |
| *LOX* | 0.08 | 0.05 | 0.097 | 0.290 | -0.09 | 0.13 | 0.502 | 0.771 |
| *LOXL1* | 0.03 | 0.06 | 0.612 | 0.832 | -0.06 | 0.17 | 0.717 | 0.807 |
| *LOXL2* | 0.01 | 0.04 | 0.857 | 0.857 | 0.06 | 0.12 | 0.643 | 0.771 |
| *SPARC* | 0.09 | 0.05 | 0.104 | 0.290 | -0.17 | 0.16 | 0.294 | 0.530 |
| *PDGFA* | 0.02 | 0.07 | 0.828 | 0.857 | 0.06 | 0.20 | 0.764 | 0.809 |
| *TGFB1* | -0.10 | 0.06 | 0.129 | 0.290 | 0.33 | 0.18 | 0.079 | 0.245 |
| *SERPINE1* | -0.03 | 0.04 | 0.424 | 0.693 | 0.17 | 0.12 | 0.179 | 0.358 |
| *CCN2* | -0.01 | 0.05 | 0.797 | 0.857 | 0.07 | 0.14 | 0.614 | 0.771 |
| *HIF1A* | -0.06 | 0.06 | 0.263 | 0.474 | 0.28 | 0.16 | 0.095 | 0.245 |
| *NLRP3* | -0.06 | 0.04 | 0.125 | 0.290 | 0.21 | 0.11 | 0.073 | 0.245 |
| *MRC1* | **-0.19** | **0.07** | **0.011** | 0.096 | **0.50** | **0.19** | **0.017** | 0.245 |
| *CD68* | -0.14 | 0.07 | 0.074 | 0.290 | 0.37 | 0.20 | 0.084 | 0.245 |

Significant relationships are in bold.

**Table 4.** Linear regression models for changes in adipose tissue fibrosis and changes in anthropometric measurements before and after bariatric surgery.

|  | SCABD Fibrosis |  |  | SCABD Fibrosis per Cell Number |  |  |
| --- | --- | --- | --- | --- | --- | --- |
|  | Estimate | Std Err | p-value | Estimate | Std Err | p-value |
| Weight (kg) | **-2.04** | **0.99** | **0.048** | -35.7 | 18.2 | 0.062 |
| BMI (kg/m<sup>2</sup>) | **-0.73** | **0.35** | **0.044** | **-14.6** | **6.36** | **0.031** |
| WC | -0.97 | 0.77 | 0.223 | -10.4 | 17.8 | 0.564 |
| HC | -1.38 | 0.79 | 0.092 | -26.3 | 16.4 | 0.126 |
| WHR | 0.01 | 0.006 | 0.875 | 0.06 | 0.14 | 0.656 |
|  | OM Fibrosis |  |  | OM Fibrosis per Cell Number |  |  |
|  | Estimate | Std Err | p-value | Estimate | Std Err | p-value |
| Weight (kg) | 0.28 | 0.93 | 0.762 | -13.9 | 24.7 | 0.579 |
| BMI (kg/m<sup>2</sup>) | 0.01 | 0.33 | 0.965 | -5.74 | 8.61 | 0.511 |
| WC | 1.33 | 0.72 | 0.076 | 17.6 | 22.4 | 0.442 |
| HC | -0.09 | 0.73 | 0.905 | 11.8 | 20.6 | 0.573 |
| WHR | 0.01 | 0.01 | 0.060 | 0.10 | 0.18 | 0.558 |

Significant relationships are in bold. SC, Subcutaneous abdominal; OM, Omental; WC, Waist Circumference; HC, Hip Circumference; WHR, Waist-to-Hip Ratio; Std Err, Standard Error.

## DISCUSSION

This study examined the effect of bariatric surgery on both SCABD and OM adipose tissue pericellular fibrosis, and on the expression of genes associated with ECM remodeling and inflammation. While mRNA expression of genes encoding for collagens in SCABD fat was not affected by weight loss, the expression of regulators of ECM remodeling significantly decreased one-year post-surgery. When measured using the usual method of picrosirius red staining, adipose tissue fibrosis was not modified two years post-surgery. Because adipocytes decrease in size following weight loss (Murphy et al., 2017), area-based measurement of adipose tissue fibrosis may overestimate the amount of collagen in samples after bariatric surgery. After expressing collagen measurements relative to the number of adipocytes, OM, but not SCABD, adipose tissue pericellular fibrosis was decreased after weight loss.

Our results contradict previous observations where the increase in pericellular collagen in SCABD fat remained significant after adjustment for adipocyte size reduction after bariatric surgery (Liu et al., 2016). On the other hand, another study found no change in SCABD fat total fibrosis 6 months post sleeve gastrectomy (Chabot et al., 2017). Likewise, after 26 weeks of liraglutide and significant weight loss, pericellular SCABD adipose tissue fibrosis remained unchanged (Wegeberg et al., 2021). It was reported that SCABD fibrosis levels were comparable between patients who experienced weight loss and those with severe obesity (BMI ≥ 40 kg/m^2^), but well above the fibrosis levels of patients with overweight although the former shared a comparable BMI (Cancello et al., 2013). It has previously been proposed that tissue with stiff ECM and advanced adipose tissue fibrosis cannot be dynamically remodeled (Lawler et al., 2016). This suggests that obesity, particularly severe obesity, may induce potentially irreversible alterations in SCABD adipose tissue ECM that are not fully reversed by substantial weight loss. Whether this applies to visceral adipose tissue is still to be determined.

We report that a decrease in SCABD adipose tissue fibrosis in patients who underwent sleeve gastrectomy was associated with greater decreases in weight and BMI. This relationship was not found for visceral fibrosis. Several studies by Clement and colleagues have demonstrated that SCABD adipose tissue fibrosis may restrict the extent of weight loss achieved following bariatric surgery (Abdennour et al., 2014; Divoux et al., 2010; Liu et al., 2016). Another study by the same group used an adipose tissue fibrosis score, which combined both perilobular and pericellular fibrosis, and found a negative relationship between said score and one year post surgery weight (Bel Lassen et al., 2017). Studies by Osorio-Conles and collaborators showed that SCABD adipose tissue mRNA levels of *COL5A1* and *COL6A3* predicted weight loss 12 months post-surgery, but no other relationship were observed between weight loss and the degree of adipose tissue fibrosis, whether assessed by histological picrosirius red staining or by biochemical measurement of hydroxyproline residues (Osorio-Conles, Jimenez, et al., 2023; Osorio-Conles, Olbeyra, et al., 2023). It is assumed that with weight changes, the ECM may adjust its support to adipocytes accordingly. However, ECM that cannot be dynamically remodeled may cause traction forces between the now smaller adipocytes and the surrounding ECM (Mariman, 2012), which may then be resolved when adipocytes return to their initial size, i.e. when weight is regained (Eastman, 2009). Another explanation may be that histological measurements of adipose tissue fibrosis simply does not adequately represents ECM remodeling. Other authors have proposed that ECM composition and dynamics, for example matrix stiffness, more than its abundance, may explain the relationship between adipose ECM and weight loss (Osorio-Conles, Olbeyra, et al., 2023; Wenderott et al., 2020).

We also investigated whether the expression of collagen-related genes may serve as markers of improvements in adipose tissue function following weight loss. We found that, while SCABD *COL5A1* mRNA expression did not change 12 months post-surgery and *COL6A3* expression tended to increase, only changes in *COL6A3* mRNA expression were negatively correlated with weight loss one year after bariatric surgery, prior to FDR adjustment. Interestingly, despite no differences in expression levels after weight loss, expression of *MMP14* also associated with weight loss resistance in the current study. The pericellular metalloproteinase MMP14 modulates pericellular collagens and is involved in overall ECM remodeling (Chun et al., 2010; Li et al., 2020). Mice with overexpression of *MMP14* in adipose tissue have increased body weight, enlarged adipocytes, and display insulin resistance (Li et al., 2020). MMP14 also cleaves COL6α3 into a subunit called endotrophin (Sun et al., 2023). In humans, plasma levels of COL6α3-derived endotrophin were strongly elevated with increasing BMI and are associated with a higher risk of coronary artery disease (Yoshiji et al., 2025) and chronic kidney disease (Staunstrup et al., 2021). Endotrophin is also a risk marker for mortality and cardiovascular complications in type 2 diabetes (Tougaard et al., 2022). Weight loss generally reduces plasma levels of COL6A3-derived endotrophin (Sachan et al., 2023; Yoshiji et al., 2025). The association between changes in adipose tissue mRNA levels in *COL6A3* and weight loss may be explained by endotrophin, although the current study may not refute or confirm this hypothesis.

We investigated the relationship previously reported between adipose tissue fibrosis and MASLD, MASH and liver fibrosis, in a population with severe obesity, and following weight loss. Unlike previous reports, we showed that in bariatric surgery candidates, adipose tissue fibrosis is a poor correlate of liver steatosis, MASH and liver fibrosis. We nonetheless detected an association between non-adjusted fibrosis in visceral adipose tissue and worsening of hepatic fibrosis following bariatric surgery-induced weight loss. Our findings contrast with those of several existing studies. It was previously shown that picrosirius red-stained fibrosis in visceral adipose tissue of patients with MASLD and obesity was significantly increased compared to a normal-weight control group, although it did not differ between patients with MASLD and MASH (Leven et al., 2021). Collagen deposition in SCABD adipose tissue was also increased in people with MASLD and obesity compared to those with obesity only (Beals et al., 2021). Both total and pericellular SCABD, but not OM, adipose tissue fibrosis associated with liver fibrosis (fibrosis score ≥ 2) in bariatric surgery patients (Abdennour et al., 2014). It was suggested that lower SCABD adipose tissue stiffness enables lipid accumulation into adipocytes, thereby limiting deposition in the liver, while increased stiffness reduces adipocyte storage capacity and promotes lipid accumulation in ectopic sites, including the liver (Virtue & Vidal-Puig, 2010). This was later challenged by Beals and colleagues, when they demonstrated that SCABD triglyceride synthesis did not differ between patients with or without MASLD, suggesting that impaired adipose tissue expandability may not be responsible for ectopic fat accumulation (Beals et al., 2021). Hepatic steatosis may rather be caused by alterations in liver physiology specifically, such as increased hepatic *de novo* lipogenesis and fatty acid uptake (Lambert et al., 2014; Smith et al., 2020). Our findings suggest that the association between MASLD features and adipose tissue fibrosis is more limited in the context of severe obesity, and adipose tissue fibrosis may not explain MASLD progression in this population.

The prevalence of MASLD is high (above 75%) in the population with obesity (Lajeunesse-Trempe et al., 2025; Quek et al., 2023). Bariatric surgery allows those patients to drastically decrease their body weight, improve glucose and lipid metabolism, while also inducing remission of comorbidities, including type 2 diabetes (Huttasch et al., 2024). Significant weight loss combined with anatomical and physiological changes associated with bariatric surgery generally leads to improvement in MASLD and liver characteristics (Fakhry et al., 2019; Hwang et al., 2024). Here, we found a small number of patients for which bariatric surgery led to aggravation or development of features associated with MASLD. Around 20% of patients are considered as non-responders to bariatric surgery and had persistent MASLD five years after surgery (Lassailly et al., 2020). This “non-response” seems to occur regardless of the type of bariatric surgery. A few studies have reported worsening fibrosis (of at least one stage) (Murakami et al., 2021; Pedersen et al., 2021) and hepatic steatosis (non-MASH turned into MASH) (Agarwal et al., 2021; Salman et al., 2020) in patients following sleeve gastrectomy. A study comparing the effect of sleeve gastrectomy and Roux-en-Y gastric bypass on MASLD noted worsening fibrosis in 12% of patients, all after Roux-en-Y gastric bypass (Froylich et al., 2016). Others have reported worsening fibrosis in patients who had little or no fibrosis preoperatively, and improvement in patients with more advanced fibrosis after biliopancreatic diversion with duodenal switch (Kral et al., 2004). However, patients with post-surgical fibrosis had other known causes of fibrosis (hemosiderosis, obstructive biliary disease, excessive alcohol consumption, reversal of their operations) leading to recurrence of metabolic syndrome. Other potential causes of liver injury occurring after bariatric surgery include aging, malnutrition, intestinal barrier dysfunction, and excessive adipose tissue lipolysis (Moolenaar et al., 2022; Salman et al., 2020). To this end, we cannot state that adipose tissue fibrosis is implicated in MASLD development and progression following weight loss, although the involvement of other adipose tissue dysfunction features cannot be excluded.

This study has many strengths, such as the use of the gold standard approach (i.e., liver biopsy) to evaluate MASLD, MASH and hepatic fibrosis, as well as the measurement of adipose tissue fibrosis from two fat compartments. However, our study included men and women of European descent, and the results may not be generalizable to other ethnic groups. The lack of a control group, such as individuals without obesity and who never had obesity, also restricts the interpretation of the results. Weight regain after bariatric surgery has several possible causes which were unexplored in this study, including elevated post-operative caloric intake, insufficient physical activity, and psychosocial stress (Noria et al., 2023). We also observed different results according to the units used to express adipose tissue fibrosis, which led to distinct or even opposite interpretations. Measurements of tissue mechanics, such as adipose tissue elasticity, may better inform on differences in adipose tissue fibrosis compared with histology-based methods (Wenderott et al., 2020). Nonetheless, we demonstrated that weight loss has distinct effects on SCABD and OM fibrosis that merit further consideration.

## Conclusion

We have demonstrated that, in severe obesity and after weight loss, histological measurement of adipose tissue fibrosis does not correlate with MASLD, MASH or hepatic fibrosis. Patients with poorer response to bariatric surgery, i.e. limited weight loss, also demonstrated less improvement in SCABD fibrosis. Weight loss nonetheless downregulates the expression of several genes involved in ECM remodeling and inflammation, indirectly suggesting a possible improvement of collagen deposition and overall adipose tissue health. These results need to be interpreted cautiously, as sometimes cofounding results were observed depending on units used to express fibrosis measurements. These findings highlight the need to reconsider methodologies and to reevaluate the clinical significance of adipose tissue fibrosis in the context of obesity and metabolic liver disease.

## Acknowledgement

The authors acknowledge the invaluable collaboration of the surgery team, bariatric surgeons and biobank staff of the QHLI.

## Statement of Ethics

This study was approved by the ethics committee of the QHLI-Laval University (approval numbers #2021–3398, #2018-2951, #2015-2466, and #2017-2710). All study-participants provided written informed consent before their inclusion in the study in compliance with the Helsinki Declaration.

## Author Contributions

EJT performed the statistical analyses, interpreted the results, and developed the draft of the manuscript. LB, SL and CYC performed bariatric surgeries on patients and/or managed clinical aspects of the study. FLT, IMP and GO were involved in adipose tissue fibrosis measurements and development of the methodological approach. MCV and JDTM performed RNA sequencing and gene expression analyses. PM and AT supervised the study and provided the scientific framework. AT also supervised the study in collaboration with the medical team. All authors read and approved the final manuscript.

## Data Availability Statement

Data used to support the findings of this study are restricted by the QHLI-Laval University (IUCPQ-UL) research ethics committee regulation to protect patient privacy. Data are available from the corresponding author (Andre Tchernof) for researchers who meet the criteria for access to confidential data.

