## Supplementary Materials for "Adipose Tissue Pericellular Fibrosis, Weight Loss and Features of Metabolic Dysfunction-Associated Steatotic Liver Disease after Bariatric Surgery"

*Research Paper*

[2]École de nutrition, Faculté des sciences de l'agriculture et de l'alimentation, Université Laval, Quebec City, QC, Canada

[3] Centre Nutrition, santé et société (NUTRISS) - Institut sur la nutrition et les aliments fonctionnels (INAF), Université Laval, Québec, QC G1V 0A6, Canada.

[4]Faculté de pharmacie, Université Laval, Quebec City, QC, Canada

[5]Département de kinésiologie, Faculté de médecine, Université Laval, Quebec City, QC, Canada

**Corresponding author:** André Tchernof, Ph.D.
Quebec Heart and Lung Institute
School of Nutrition, Université Laval
2725 Chemin Sainte-Foy (Y-4212)
Quebec City, QC
CANADA G1V 4G5


**Keywords**: Fibrosis, Picrosirius Red, Severe Obesity, Bariatric and Metabolic Surgery, MASLD, MASH, Liver Fibrosis

**Fibrose – Supplementary Tables**

**Supplementary Table 1**: Transcripts examined in this study and their functions

| **Genes** | **Functions** | **References** |
|---|---|---|
| Collagen Type V Alpha 1 Chain (*COL5A1*) | Regulatory fibril-forming collagen | (Mak et al., 2016) |
| Collagen Type VI Alpha 3 Chain (*COL6A3*) | Beaded filament collagen | (Khan et al., 2009; Pasarica et al., 2009) |
| Lysyl Oxidase (*LOX*, *LOX1*, *LOX2*) | Collagen stability | (Pastel et al., 2018) |
| Metalloproteinase (*MMP2*, *MMP9*, *MMP14*) and Metalloproteinase Inhibitor (*TIMP2*) | Extracellular matrix proteins degradation, and their inhibitors | (Berg et al., 2014; Sachan et al., 2023) |
| Transforming Growth Factor Beta 1 (*TGFB1*) | Promotes fibronectin and collagen production | (Fain et al., 2005; Meng et al., 2016) |
| Platelet-Derived Growth Factor Subunit A (*PDGFA*) | Fibrogenic cytokine | (Marcelin et al., 2017) |
| Connective Tissue Growth Factor (*CCN2*) | Pro-fibrotic factor | (Yoshino et al., 2019) |
| Endothelial Plasminogen Activator Inhibitor (*SERPINE1*) | Fibrinolysis inhibitor | (Kaji, 2016) |
| Secreted Protein Acidic and Rich in Cysteine (*SPARC*) | Regulates cell growth through interactions with the extracellular matrix | (Kos & Wilding, 2010) |
| Macrophage Markers Cluster of Differentiation 68 (*CD68*) | Marker of macrophages and monocytes | (Engin, 2017; Moreno-Navarrete et al., 2013) |
| Mannose Receptor C-Type 1 (*MRC1*) | Marker of macrophages, dendritic cells, and endothelial cells | (Engin, 2017; Moreno-Navarrete et al., 2013) |
| Hypoxia-Inducible Factor 1-Alpha (*HIF1A*) | Regulator of cellular and systemic homeostatic response to hypoxia | (Halberg et al., 2009; Sun et al., 2013) |
| NOD-Like Receptor Family Pyrin Domain Containing 3 (*NLRP3*) | Regulator of inflammation | (Pahwa et al., 2021) |

**Supplementary Table 2**: Number and proportion of participants with MASLD, MASH and hepatic fibrosis before and after bariatric surgery (patient sample 3)

| **Histopathological characteristics** | **First surgery n (%)** | **Second surgery (26 ± 6 months) n (%)** | **p-value** |
|---|---|---|---|
| **Steatosis** | | | |
| Grade 0 | 4 (10.8) | 17 (47.2) | 0.001 |
| Grade 1 | 21 (56.8) | 17 (47.2) | |
| Grade 2 | 8 (21.6) | 2 (5.6) | |
| Grade 3 | 4 (10.8) | 0 (0) | |
| **MASH** | | | |
| No | 20 (54.0) | 30 (81.1) | 0.013 |
| Yes | 17 (46.0) | 7 (18.9) | |
| **Hepatic Fibrosis** | | | |
| Grade 0 | 17 (46.0) | 25 (67.6) | 0.173 |
| Grade 1 | 10 (27.0) | 4 (10.8) | |
| Grade 2 | 8 (21.6) | 5 (13.5) | |
| Grade 3 | 2 (5.4) | 3 (8.1) | |

MASLD, Metabolic dysfunction–associated steatotic liver disease; MASH, metabolic dysfunction-associated steatohepatitis.

**Supplementary Table 3**: Number and proportion of patients with maintained, improved, unchanged or worsened hepatic characteristics after bariatric surgery (patient sample 3)

| **Characteristics** | **Maintained** | **Improved** | **Unimproved** | **Worsened** | ***De novo*** |
|---|---|---|---|---|---|
| Steatosis, n (%) | 3 (8.6) | 23 (65.7) | 8 (22.9) | 1 (2.8) | - |
| MASH, n (%) | 16 (44.5) | 13 (36.1) | 4 (11.1) | - | 3 (8.3) |
| Hepatic Fibrosis, n (%) | 11 (30.6) | 12 (33.3) | 7 (19.4) | 2 (5.6) | 4 (11.1) |

**Fibrose – Supplementary Figures**

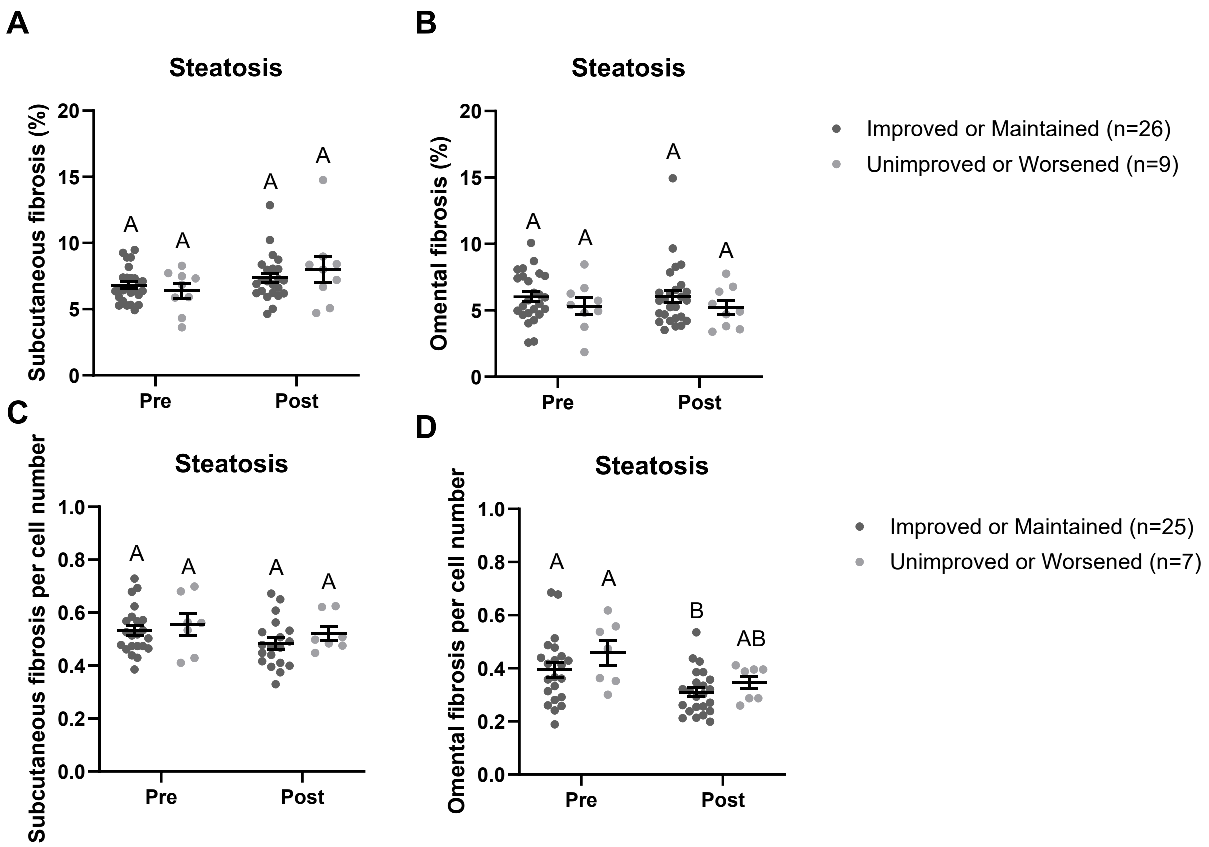


**Supplementary Figure 1**. Adipose tissue fibrosis measured by picrosirius red labeling of collagen expressed as area (A, B) or area by cell number (C, D) in two fat compartments from patients with improved/maintained or unimproved/worsened steatosis grade after bariatric surgery. P-values are adjusted for age and sex. ANOVA and Tukey, means ± SEM, groups showing different letters are statistically different from each other at $p<0.05$.

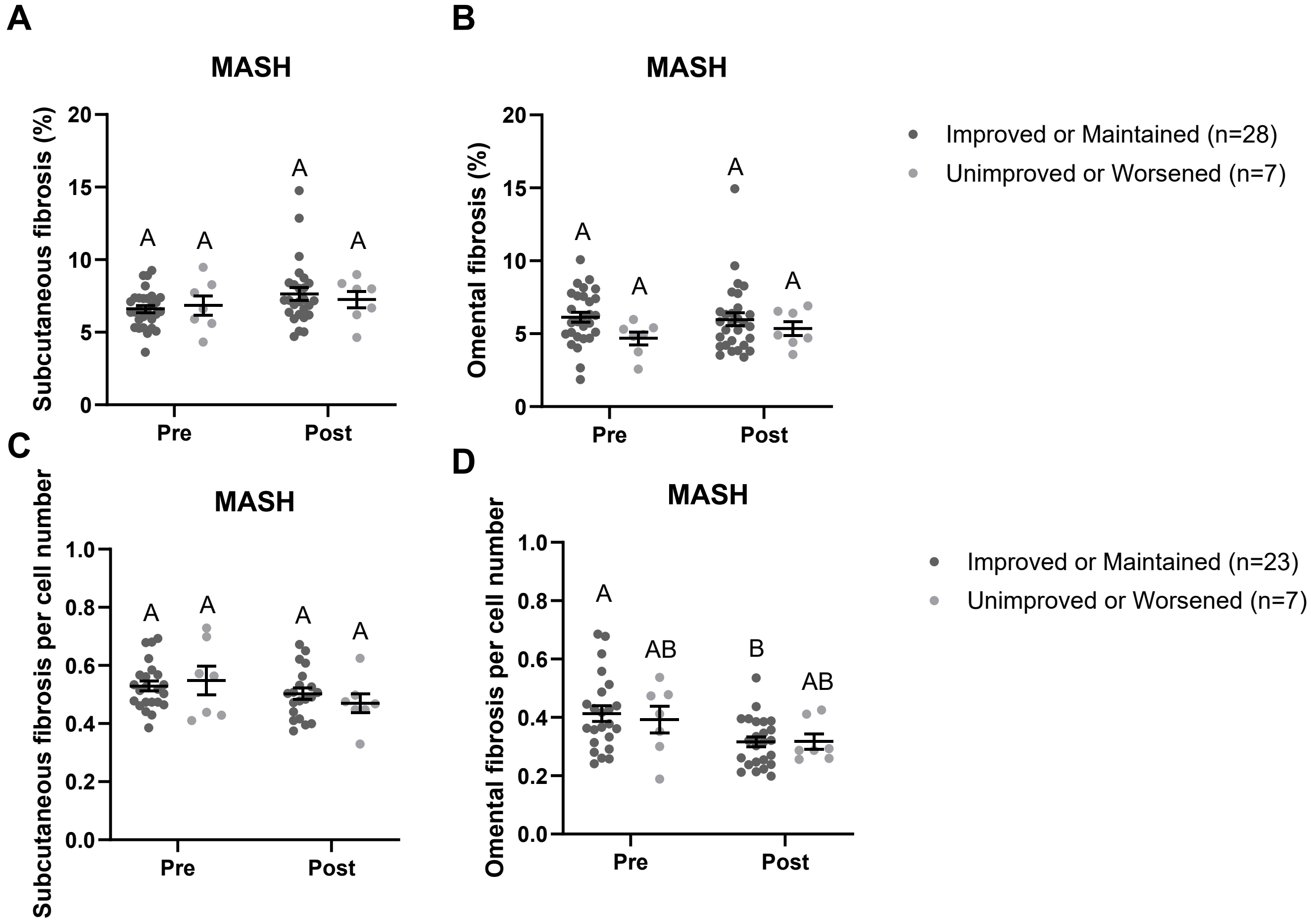


**Supplementary Figure 2**. Adipose tissue fibrosis measured by picrosirius red labeling of collagen expressed as area (A, B) or area by cell number (C, D) in two fat compartments from patients with improved/maintained or unimproved/worsened MASH after bariatric surgery. P-values are adjusted for age and sex. ANOVA and Tukey, means ± SEM, groups showing different letters are statistically different from each other at $p<0.05$.

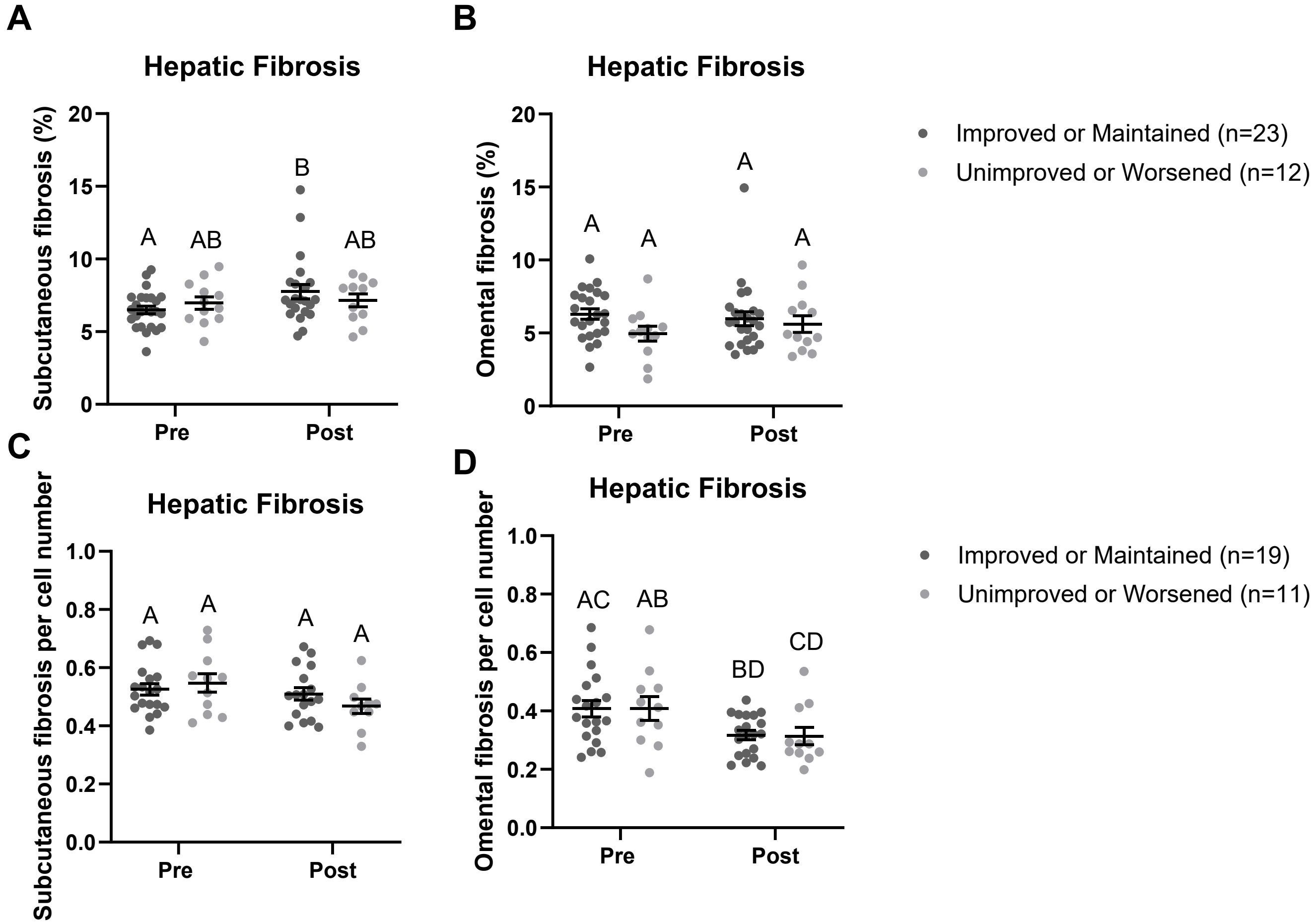


**Supplementary Figure 3**. Adipose tissue fibrosis measured by picrosirius red labeling of collagen expressed as area (A, B) or area by cell number (C, D) in two fat compartments from patients with improved/maintained or unimproved/worsened hepatic fibrosis grade after bariatric surgery. P-values are adjusted for age and sex. ANOVA and Tukey, means ± SEM, groups showing different letters are statistically different from each other at $p<0.05$.

Berg, G., Schreier, L., & Miksztowicz, V. (2014). Circulating and adipose tissue matrix metalloproteinases in cardiometabolic risk environments: pathophysiological aspects. *Horm Mol Biol Clin Investig*, *17*(2), 79-87. https://doi.org/10.1515/hmbci-2013-0069

Engin, A. B. (2017). Adipocyte-Macrophage Cross-Talk in Obesity. *Adv Exp Med Biol*, *960*, 327-343. https://doi.org/10.1007/978-3-319-48382-5_14

Fain, J. N., Tichansky, D. S., & Madan, A. K. (2005). Transforming growth factor beta1 release by human adipose tissue is enhanced in obesity. *Metabolism*, *54*(11), 1546-1551. https://doi.org/10.1016/j.metabol.2005.05.024

Halberg, N., Khan, T., Trujillo, M. E., Wernstedt-Asterholm, I., Attie, A. D., Sherwani, S., Wang, Z. V., Landskroner-Eiger, S., Dineen, S., Magalang, U. J., Brekken, R. A., & Scherer, P. E. (2009). Hypoxia-inducible factor 1alpha induces fibrosis and insulin resistance in white adipose tissue. *Mol Cell Biol*, *29*(16), 4467-4483. https://doi.org/10.1128/MCB.00192-09

Kaji, H. (2016). Adipose Tissue-Derived Plasminogen Activator Inhibitor-1 Function and Regulation. *Compr Physiol*, *6*(4), 1873-1896. https://doi.org/10.1002/cphy.c160004

Khan, T., Muise, E. S., Iyengar, P., Wang, Z. V., Chandalia, M., Abate, N., Zhang, B. B., Bonaldo, P., Chua, S., & Scherer, P. E. (2009). Metabolic dysregulation and adipose tissue fibrosis: role of collagen VI. *Mol Cell Biol*, *29*(6), 1575-1591. https://doi.org/10.1128/MCB.01300-08

Kos, K., & Wilding, J. P. (2010). SPARC: a key player in the pathologies associated with obesity and diabetes. *Nat Rev Endocrinol*, *6*(4), 225-235. https://doi.org/10.1038/nrendo.2010.18

Mak, K. M., Png, C. Y., & Lee, D. J. (2016). Type V Collagen in Health, Disease, and Fibrosis. *Anat Rec (Hoboken)*, *299*(5), 613-629. https://doi.org/10.1002/ar.23330

Marcelin, G., Ferreira, A., Liu, Y., Atlan, M., Aron-Wisnewsky, J., Pelloux, V., Botbol, Y., Ambrosini, M., Fradet, M., Rouault, C., Henegar, C., Hulot, J. S., Poitou, C., Torcivia, A., Nail-Barthelemy, R., Bichet, J. C., Gautier, E. L., & Clement, K. (2017). A PDGFRalpha-Mediated Switch toward CD9(high) Adipocyte Progenitors Controls Obesity-Induced Adipose Tissue Fibrosis. *Cell Metab*, *25*(3), 673-685. https://doi.org/10.1016/j.cmet.2017.01.010

Meng, X. M., Nikolic-Paterson, D. J., & Lan, H. Y. (2016). TGF-beta: the master regulator of fibrosis. *Nat Rev Nephrol*, *12*(6), 325-338. https://doi.org/10.1038/nrneph.2016.48

Moreno-Navarrete, J. M., Ortega, F., Gomez-Serrano, M., Garcia-Santos, E., Ricart, W., Tinahones, F., Mingrone, G., Peral, B., & Fernandez-Real, J. M. (2013). The MRC1/CD68 ratio is positively associated with adipose tissue lipogenesis and with muscle mitochondrial gene expression in humans. *PLoS One*, *8*(8), e70810. https://doi.org/10.1371/journal.pone.0070810

Pahwa, R., Singh, A., Adams-Huet, B., Devaraj, S., & Jialal, I. (2021). Increased inflammasome activity in subcutaneous adipose tissue of patients with metabolic syndrome. *Diabetes Metab Res Rev*, *37*(3), e3383. https://doi.org/10.1002/dmrr.3383

Pasarica, M., Gowronska-Kozak, B., Burk, D., Remedios, I., Hymel, D., Gimble, J., Ravussin, E., Bray, G. A., & Smith, S. R. (2009). Adipose tissue collagen VI in obesity. *J Clin Endocrinol Metab*, *94*(12), 5155-5162. https://doi.org/10.1210/jc.2009-0947

Pastel, E., Price, E., Sjoholm, K., McCulloch, L. J., Rittig, N., Liversedge, N., Knight, B., Moller, N., Svensson, P. A., & Kos, K. (2018). Lysyl oxidase and adipose tissue dysfunction. *Metabolism*, *78*, 118-127. https://doi.org/10.1016/j.metabol.2017.10.002

Sachan, A., Aggarwal, S., Pol, M. M., Singh, A., & Yadav, R. (2023). Expression analysis of MMP14: Key enzyme action in modulating visceral adipose tissue plasticity in patients with obesity. *Clin Obes*, *13*(5), e12607. https://doi.org/10.1111/cob.12607

Sun, K., Halberg, N., Khan, M., Magalang, U. J., & Scherer, P. E. (2013). Selective inhibition of hypoxia-inducible factor 1alpha ameliorates adipose tissue dysfunction. *Mol Cell Biol*, *33*(5), 904-917. https://doi.org/10.1128/MCB.00951-12

Yoshino, J., Patterson, B. W., & Klein, S. (2019). Adipose Tissue CTGF Expression is Associated with Adiposity and Insulin Resistance in Humans. *Obesity (Silver Spring)*, *27*(6), 957-962. https://doi.org/10.1002/oby.22463
